# Recent genetic drift in the co-diversified gut bacterial symbionts of laboratory mice

**DOI:** 10.1101/2024.08.14.607958

**Authors:** Daniel D. Sprockett, Brian A. Dillard, Abigail A. Landers, Jon G. Sanders, Andrew H. Moeller

**Affiliations:** Department of Ecology and Evolutionary Biology, Cornell University, Ithaca, NY 14853, USA; Department of Ecology and Evolutionary Biology, Princeton University, Princeton, NJ 08540, USA

## Abstract

Laboratory mice (*Mus musculus domesticus*) harbor gut bacterial strains that are distinct from those of wild mice^1^ but whose evolutionary histories are poorly understood. Understanding the divergence of laboratory-mouse gut microbiota (LGM) from wild-mouse gut microbiota (WGM) is critical, because LGM and WGM have been previously shown to differentially affect mouse immune-cell proliferation^2,3^, infection resistance^4^, cancer progression^2^, and ability to model drug outcomes for humans^5^. Here, we show that laboratory mice have retained gut bacterial symbiont lineages that diversified in parallel (co-diversified) with rodent species for > 25 million years, but that LGM strains of these ancestral symbionts have experienced accelerated accumulation of genetic load during the past ∼ 120 years of captivity. Compared to closely related WGM strains, co-diversified LGM strains displayed significantly faster genome-wide rates of fixation of nonsynonymous mutations, indicating elevated genetic drift, a difference that was absent in non-co-diversified symbiont clades. Competition experiments in germ-free mice further indicated that LGM strains within co-diversified clades displayed significantly reduced fitness *in vivo* compared to WGM relatives to an extent not observed within non-co-diversified clades. Thus, stochastic processes (*e.g.*, bottlenecks), not natural selection in the laboratory, have been the predominant evolutionary forces underlying divergence of co-diversified symbiont strains between laboratory and wild house mice. Our results show that gut bacterial lineages conserved in diverse rodent species have acquired novel mutational burdens in laboratory mice, providing an evolutionary rationale for restoring laboratory mice with wild gut bacterial strain diversity.

---

Mammalian lineages share gut bacterial taxa (*e.g.*, genera), but bacterial strains within these taxa differ systematically among host populations and species^6–12^. For example, common laboratory lines of house mice (*Mus musculus domesticus*) (*e.g.*, C57BL/6J), which were derived from wild mice > 100 years ago^13^, harbor gut-microbiota strains distinct from those of the same taxa found within wild house mice^1^. The evolutionary histories of laboratory mouse–gut microbiota (LGM) strains remain unclear. LGM strains may have been acquired in captivity from external sources, such as humans or the indoor environment. Alternatively, LGM strains may be descendants of ancestral symbionts from the wild-mouse gut microbiota (WGM) that have been reshaped by laboratory-specific evolutionary forces, such as natural selection or genetic drift.

In principle, LGM strains descended from symbioses ancestral to laboratory and wild mice could be identified through co-phylogenetic analyses, which test for parallel diversification (co-diversification) between symbionts and hosts. Discovering ancestral, co-diversifying symbionts in the gut-microbiota (GM) would also provide a phylogenetic framework for interrogation of the histories of symbiont adaptation and genetic drift within host lineages, as has been possible in simpler host-microbe symbioses, such as those between insects and bacterial endosymbionts^14,15^. However, co-phylogenetic tests have not been feasible for rodent gut microbiota due to the limited phylogenetic information provided by 16S rDNA sequencing and assembly-free shotgun metagenomics^16^. Analyses of high-quality bacterial genomes assembled directly from metagenomes^17,18^ provide increased power to test for co-diversification between gut microbiota and hosts. These approaches have recently revealed that multiple GM strains have co-diversified with humans and other primate species^8,10^ but have yet to be leveraged to assess the evolutionary histories of mouse GM strains, owing to a lack of genome-resolved datasets from closely related rodent species.

## A genome-resolved bacterial phylogeny from the rodent gut microbiota

Here, we employed comparative, genome-resolved metagenomics approaches to test for co-diversification between GM strains and rodents, to assess the extent to which ancestral GM symbionts have persisted within laboratory mice, and to test how natural selection and genetic drift have driven divergence of LGM strains from their wild relatives. To enable tests for co-diversification between gut microbiota and rodents in the super family Muroidea, we used a combination of long-read Oxford Nanopore and short-read Illumina sequencing to generate 504 metagenome-assembled genomes (MAGs) from individuals of six species/subspecies of deer mice (genus *Peromyscus*) descended from wild populations in the United States and reared in a common facility on a common diet (Methods; Extended Data Table 1). We combined these MAGs with previously published MAGs from the gut microbiota of 307 host individuals from 14 rodent species^1,2,5,19–28^ (Fig. 1a; Extended Data Table 2) and generated a curated host tree from timetree.org^29^. We used IQTree2^30^ to construct a maximum likelihood phylogenetic tree of all MAGs based on single-copy core genes (Methods), which define bacterial lineages’ taxonomic classifications^31^. The final tree included 5,567 MAGs from 20 rodent host species, allowing assessment of GM-strain diversification coincident with rodent speciation.

**Fig. 1:**
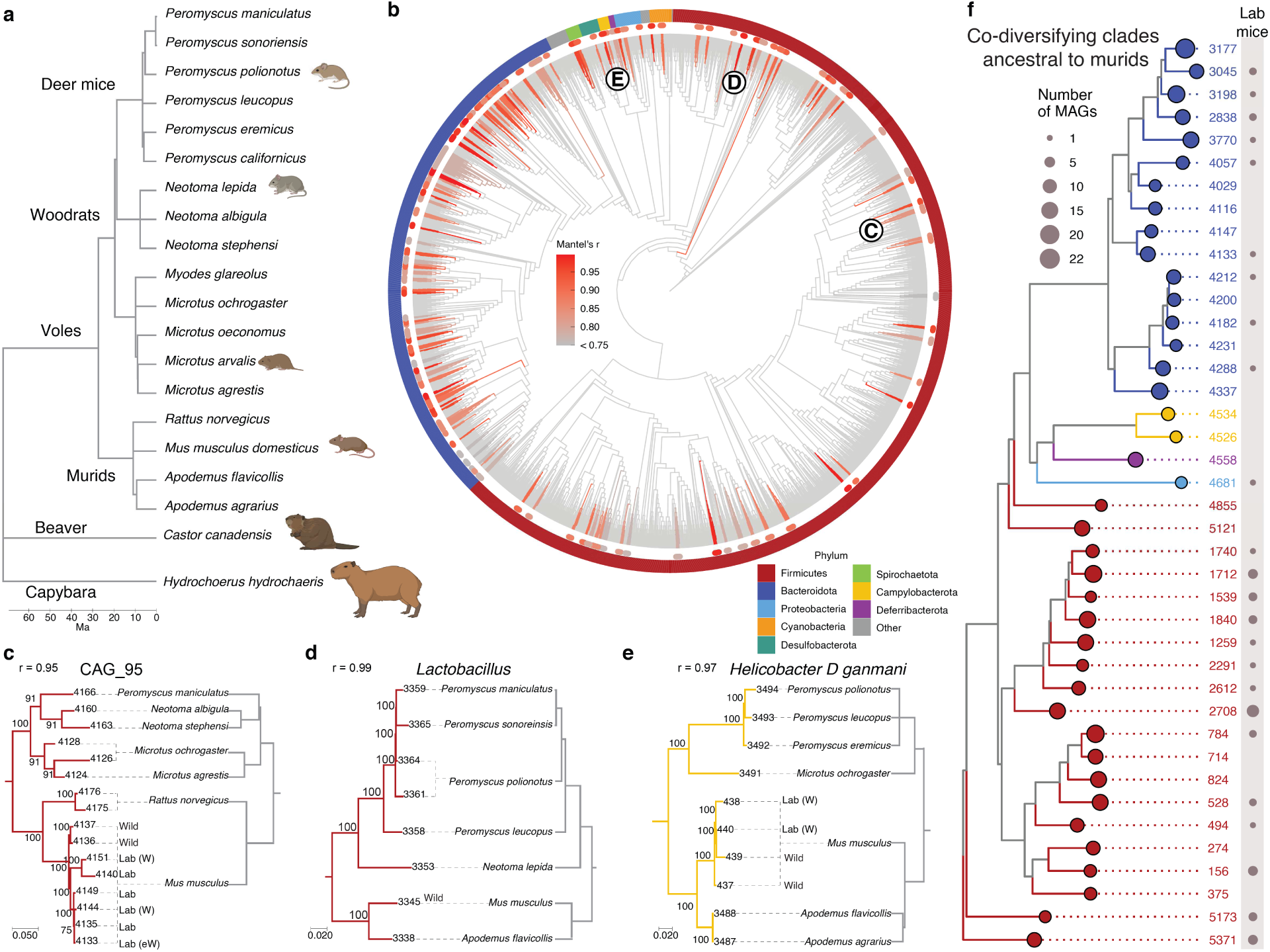
Retention of ancestral, co-diversifying rodent gut symbionts in laboratory mice. **a**, Phylogeny shows evolutionary relationships among rodent host species for which gut bacterial genomes were included in this study. **b,** Phylogeny shows relationships among gut bacterial genomes assembled from the rodent gut microbiota. Colors denote bacterial phyla. Bars surrounding phylogeny mark clades displaying significant evidence of co-diversification (r > 0.75, *p*-value < 0.01). **c–e,** Tanglegrams show concordance between gut bacterial (left) and rodent (right) phylogenetic trees. Dashed lines connect bacterial genomes with the host species from which they were recovered. Genomes derived from mice (*Mus musculus*) sampled in the lab or the wild are labeled. ‘W’ and ‘eW’ indicate ‘wildling’ and ‘ex-wild’ mice, respectively. Numbers left and right of tree are bootstrap support values and genome IDs, respectively. **f,** Phylogeny shows relationships among all bacterial genomes belonging to co-diversifying clades shown in Fig. 1 that contain genomes from murids and non-murids, *i.e.*, co-diversifying clades that could be inferred to be ancestral to murids. Circles indicate clades (colored by phylum, left) or MAGs derived from laboratory mice (grey, right). Sizes of circles indicate the number of MAGs. Identification numbers are listed to the right of each clade.

## Widespread co-diversification between gut bacterial symbionts and rodents

To test for co-diversification between GM strains and rodent species, we employed an approach previously used to identify GM strains that co-diversified with primate species^10^. We tested every node in the distal 1/4^th^ of the bacterial phylogeny (1,245 nodes) for co-diversification with host species using the method developed by Hommola *et al.*^32^ (Methods), since previous molecular clock analyses suggest that more basal nodes likely predate the most-recent common ancestors of rodents^10^. We identified 158 gut bacterial clades showing evidence of co-diversification with host species following previously employed significance thresholds of r > 0.75 and *p*-value < 0.01 (Extended Data Table 3, Fig. 1b)^10^. Co-diversifying gut bacterial lineages comprised 22.6% of the total branch length on the MAG phylogeny and spanned 8 phyla. Perfect or near perfect concordance between host and symbiont phylogenies was observed, indicative of ancestral relationships between host lineages and host species–specific symbionts spanning > 25 million years (Fig. 1c–e, Extended Data Figure 1). These include strains of *Helicobacter ganmani* (a common commensal of laboratory mice^33^), an unclassified species of *Lactobacillus* (paralleling results from studies of this taxon in other hosts^34^), several unclassified genera, and others (Extended Data Table 3). Interestingly, within these co-diversifying clades, MAGs derived from laboratory mice formed sister clades to MAGs derived from wild mice (*e.g.*, Fig. 1 c, e), ‘wildlings’^5^ (laboratory mice born to a wild-caught mother via embryo transfer), and ‘ex-wild’ mice (wild-caught mice housed in the laboratory animal facilities) (Extended Data Table 3). The phylogenetic relationships among LGM and WGM strains indicate that LGM strains are descended from ancestral symbiont lineages that resided within wild house mice and the common ancestors of wild house mice and other Muroidea species.

To further assess the evidence for co-diversification in these clades, we conducted tests after subsampling a single MAG from each monophyletic clade derived from a single host species. These analyses assessed only the association between the backbone of each symbiont clade and the host phylogeny, eliminating the possibility of pseudoreplication caused by repeated sampling of individual bacterial clades from the same host species^35^. We employed multiple tests for co-diversification, including PACo^36^, ParaFit^37^, and Hommola’s test^32^. 324 clades displaying significant evidence of co-diversification in at least one test (Extended Data Table 3, Extended Data Figure 2), the results of these different tests were significantly associated with one another (Extended Data Figure 2A–C), and 156 clades displayed significant evidence for co-diversification based on at least two of the tests (Extended Data Figure 2D). We observed between eight-fold and twelve-fold more significantly co-diversifying clades (based on a *p*-value of 0.01) than expected under the null hypothesis (*i.e.*, 1% of tests) (Extended Data Figure 3), depending on the specific test employed. In addition, we assessed the false discovery rate of the initial scan for co-diversification based on the complete dataset using a previously developed permutation test^10,38^, finding that the scan based on the real data detected always detected more co-diversifying clades than the number detected in the permuted scans, which averaged 53.74 clades (Extended Data Figure 3). Sensitivity analyses further indicated that the detection of 73– 100% of co-diversifying clades was robust to the removal of MAGs from individual host species (depending on the host species removed) (Extended Data Figure 4), and molecular clock analyses corroborated contemporaneous diversification of symbiont and rodent lineages (Extended Data Figure 5, Extended Data Table 4). Cumulatively, these results demonstrate widespread co-diversification between gut microbiota and rodent species.

## Retention and extinction of co-diversifying symbionts in laboratory house mice

The discovery of co-diversifying GM strains enabled identification of ancestral GM lineages that have either been retained in or lost from laboratory house mice. Of the 158 co-diversifying clades identified, 40 phylogenetically independent (*i.e.*, non-nested) clades were inferred to be ancestral to house mice and other murids (*i.e.*, present in *Mus musculus domesticus*, at least one non-*M. m. domesticus* murid, and at least one outgroup to murids) (Supplementary Information). Of these 40 ancestral clades, 24 contained MAGs from laboratory house mice (Fig. 1f, Extended Data Figure 6). Previous work showed that wild-derived inbred mouse lines could retain subsets of the microbiota from their wild population of origin for > 10 host generations^39^. The observation of 24 ancestral, co-diversifying clades in laboratory mice (Fig. 1f) shows that these symbioses have persisted since their hosts’ derivation from wild stock > 100 years ago^13^.

Our analyses also indicated that laboratory house mice have experienced elevated rates of loss of ancestral symbionts relative to wild house mice. Only 7 ancestral clades lacked MAGs from wild house mice, whereas 33 clades contained MAGs from wild house mice (compared to 16 and 24 clades, respectively, for laboratory house mice). These results indicate significantly greater absence of these clades from laboratory house mice than from wild house mice (chi-squared test *p*-value = 0.0262), consistent with accelerated loss of ancestral gut-microbiota diversity from laboratory house mice^1^. This difference could not be explained by sampling effort, which favored laboratory house mice (217 laboratory versus 90 wild house-mouse samples).

MAGs from ‘wildling’^5^ or ‘rewilded’^3,4^ mice (lab mice released outdoors) were contained within a subset of clades lacking MAGs from laboratory mice (Extended Data Figure 6), indicating that ancestral clades absent from laboratory mice can be regained through ‘wildling’ or ‘rewilding’ approaches.

## Altered genomic signatures of positive selection in LGM strains

Within co-diversifying clades ancestral to murids (Fig. 1f), LGM and WGM strains repeatedly formed reciprocally monophyletic clades (*e.g.*, Fig. 1c, e), indicating their genomic distinctiveness. We next interrogated the evolutionary forces that have driven this divergence. We reasoned that LGM strains may be experiencing altered forces of natural selection compared to their wild relatives due to inbred host genetic backgrounds, laboratory-mouse diets, and myriad other factors. To test this idea, we conducted genome-wide scans for positive selection on each protein-coding gene in each co-diversifying clade ancestral to murids that contained MAGs from laboratory and wild house mice. For each gene, we calculated the ratio of the rates of nonsynonymous and synonymous substitutions per site, *i.e.*, dN/dS, which is expected to be 1 under neutral evolution, >1 under positive selection to change the protein product, and < 1 under purifying selection against non-synonymous mutations. Results showed that most co-diversifying GM strains’ genes have evolved under purifying selection in both wild and laboratory mice (Fig. 2, Extended Data Table 5), as expected. However, distinct sets of genes exhibited evidence of positive selection in LGM and WGM strains (Fig. 2). Significantly more genes displayed dN/dS > 1 in laboratory mice (76 genes) but dN/dS < 1 in wild mice than dN/dS > 1 in wild mice but dN/dS < 1 in laboratory mice (50 genes) (binomial test p-value = 0.0255), consistent with novel selective forces driving accelerated evolution of a minority of protein sequences encoded by the genomes of LGM strains. Genes showing evidence of positive selection in the laboratory but purifying selection (or near neutrality) in the wild included *hpt* (Borkfalkiaceae) involved in the purine salvage pathway, *tsaD* (Muribaculaceae) involved in tRNA metabolism, *oppC* (*Schaedlerella*) involved in oligopeptide transport, *kbl* (*Odoribacter*) involved in amino acid degradation, *carB_2* (*Duncaniella freteri*) involved in pyrimidine metabolism, *pgi* (unclassified genus 1XD8-76) involved in carbohydrate degradation, and *pflB* (*Lachnospira*) involved in pyruvate fermentation (Fig. 2; Extended Data Table 5). Laboratory-specific positive selection on these metabolic genes may result from the compositionally distinct, *ad libitum* diets of laboratory mice, although our analyses were not able to identify specific selective agents. We also tested for differences in gene functional annotations between laboratory and wild GM strains (and co-diversifying and non-co-diversifying clades, Supplementary Information), but the left-skewed distributions of p-values obtained indicated that these analyses lacked power (Extended Data Table 6, S7, respectively), suggesting relatively greater applicability of sequence-based scans for selection (*e.g.*, dN/dS) for MAGs. These results identify genes in ancestral, co-diversifying symbionts of house mice showing evidence of laboratory-specific adaptation.

**Fig. 2:**
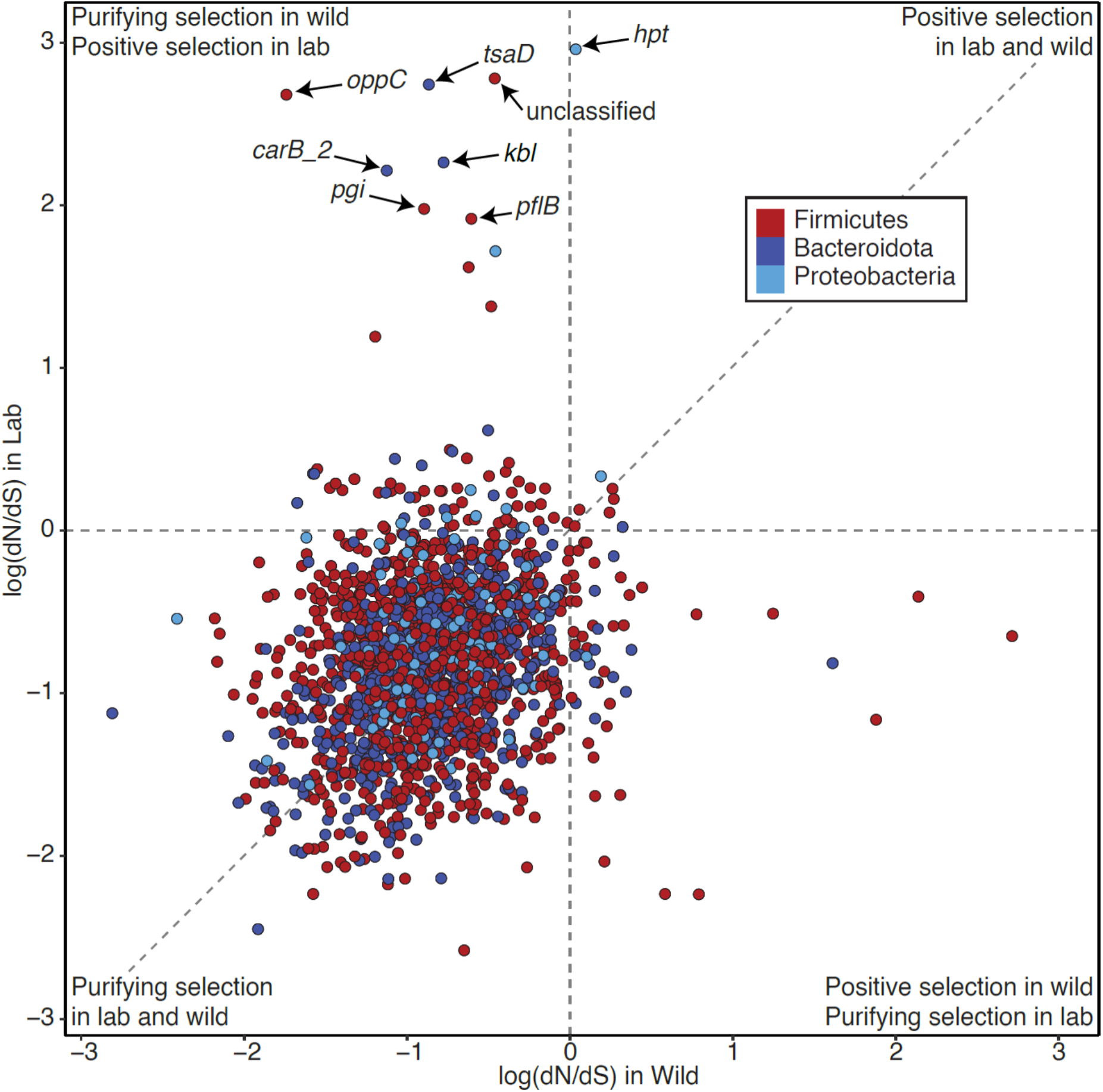
Divergent positive selection on ancestral GM symbiont strains in wild and laboratory mice. Scatter plot shows the log of the dN/dS ratio for bacterial genes in co-diversifying clades in laboratory house mice (y-axis) and other rodents (x-axis). Each point represents a bacterial gene. Positive values indicate evidence of positive selection, values near zero indicate neutral evolution, and negative values indicate purifying/negative selection. Points in the upper left quadrant show evidence of positive selection in laboratory house mice but purifying/negative selection in other rodents.

## Significantly elevated genetic drift in LGM strains

In addition to testing for divergent natural selection between LGM and WGM strains, we also tested for divergence in the strength of genetic drift. Laboratory breeding and animal care procedures may exert bottlenecks on LGM strains, which would elevate the strength of genetic drift. C57BL/6J—the most widely used laboratory mouse line—was derived from a single mating pair (https://www.jax.org/strain/000664), and previous work has shown that moving wild mice into the laboratory and establishing inbred lines is associated with a precipitous loss of GM diversity^39^, a hallmark of drift. Moreover, once in the laboratory, GM strains are vertically inherited from mother to offspring within inbred mouse lines^39^ through stochastic sampling processes that can exert bottlenecks on GM diversity^40^.

Theory predicts that stronger genetic drift will reduce the efficacy of purifying selection, increasing the rate of fixation of weakly deleterious mutations. In bacterial lineages with low effective population sizes (*N_e_*), stochastic sampling leads to elevated genome-wide rates of fixation of nonsynonymous substitutions (dN) that would otherwise be efficiently purged by purifying selection in lineages with large *N_e,_* elevating dN/dS genome-wide^41,42^. Whereas elevated dN/dS values in individual genes can be driven by the action of positive selection, elevated genome-wide dN/dS is indicative of reduced *N_e_* and increased genetic drift^43^.

To test whether co-diversifying GM strains in laboratory mice have experienced elevated genetic drift, we compared genome-wide dN/dS values in LGM and WGM strains within each co-diversifying clade ancestral to murids. We observed significant genome-wide elevation of dN/dS in LGM strains in tests based on all genes (paired-test *p*-value = 0.000356) and in tests based only on genes under purifying selection (bottom-left quadrant of Fig. 2) (paired t-test *p-* value = 0.00511) (Fig. 3a). The genome-wide signatures of genetic drift became more evident when individual co-diversifying clades were tested separately (Fig. 3b–d). Significantly elevated dN/dS in LGM strains was observed in clades of *Schaedlerella* (class: Clostridia) (paired t-test FDR-corrected *p-*values = 2.05e-07, all genes; 5.87e-06, genes under purifying selection) (Fig. 3b), unclassified Lachnospiraceae genus CAG-95 (paired t-test FDR-corrected *p-*values < 5.5e-15) (Fig. 3c), *Anaerotingum* (class: Clostridia) (paired t-test FDR-corrected *p-*values = 0.00390 and 0.0228) (Fig. 3d), and others (Extended Data Table 5). Only one co-diversifying clade, unclassified Oscillospiraceae UMGS1872, displayed significantly higher dN/dS in the wild than in the laboratory (paired t-test *p-*value = 7.93e-05 and 4.51e-05). Unclassified Oscillospiraceae were previously shown to be two of seven bacterial taxa found at significantly higher frequency in laboratory mice than in wild mice (compared to 68 taxa found at significantly higher frequency in wild mice)^1^. Larger census populations sizes of this taxon may buffer against stronger genetic drift in the laboratory. Elevated genetic drift in LGM strains was not observed for non-co-diversifying symbiont clades (Mantel’s r < 0) (Supplementary Information; Extended Data Table 8), indicating that elevated genetic drift in the laboratory has primarily affected co-diversifying bacterial symbionts. Moreover, genome-wide dN/dS was significantly higher for co-diversified clades (r > 0.75) than for non-co-diversified clades (r < 0) in both laboratory and wild house mice (Supplementary Information, Extended Data Figure 7), further indicating stronger genetic drift in co-diversifying compared to non-co-diversifying GM symbiont lineages.

**Fig. 3:**
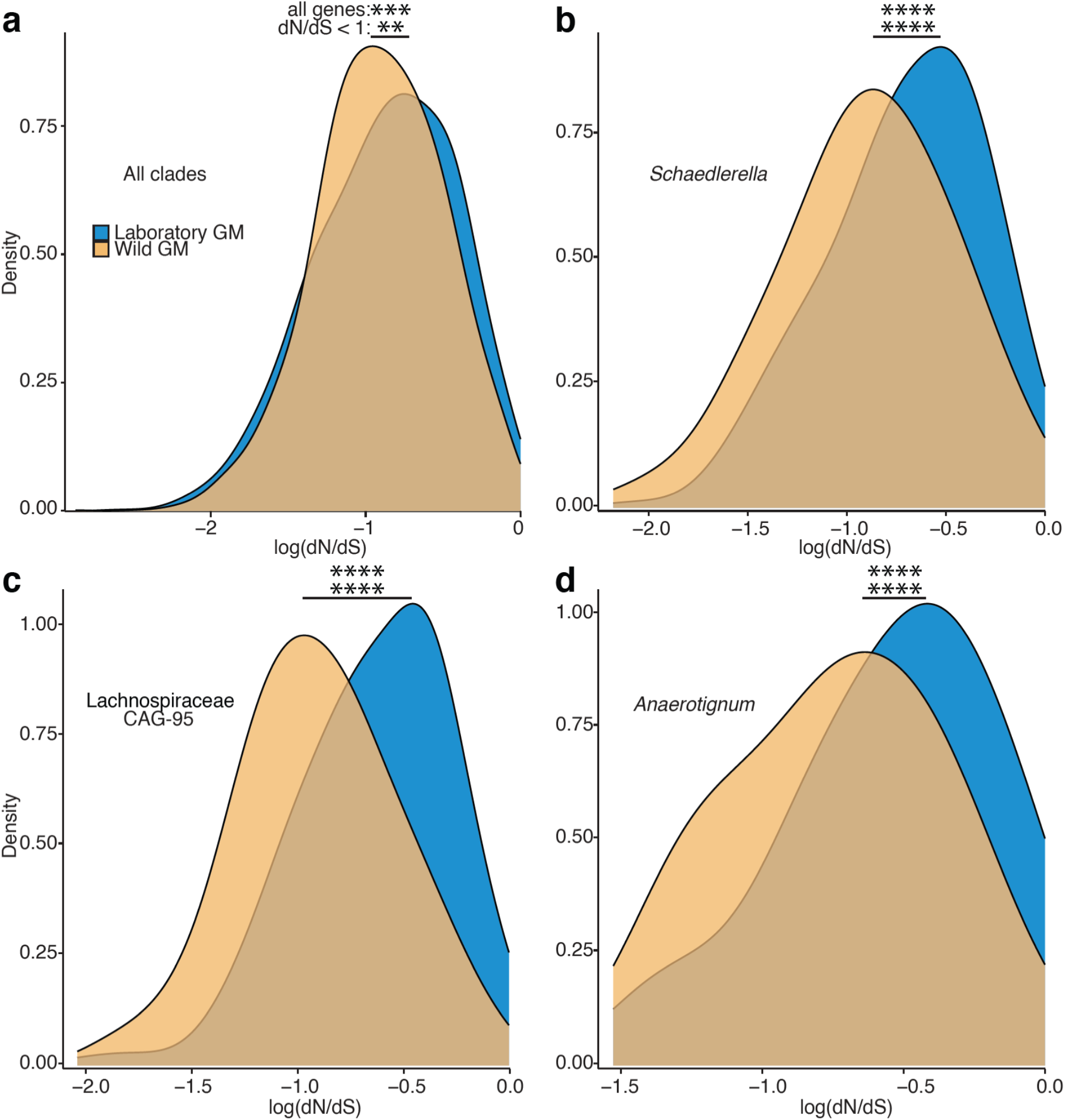
Elevated mutational load in co-diversified LGM strains compared to WGM relatives. **a**, Density plot shows significant genome-wide elevation of dN/dS (a hallmark of genetic drift) in co-diversifying GM strains in laboratory house mice relative to wild house mice. dN/dS values for all genes from all co-diversifying clades ancestral to murids and containing MAGs from both laboratory and wild mice are shown. **b–d**, Density plots show significant genome-wide elevation of dN/dS in individual co-diversifying clades in laboratory house mice relative to wild house mice. In **a–d**, significance of paired t-tests for differences in mean is denoted by asterisks; ** *p*-value < 0.01; *** *p*-value < 0.001; **** *p*-value < 0.0001. Top asterisks denote significance of tests considering all genes, and bottom asterisks denote significance of tests considering only genes displaying dN/dS < 1 (log dN/dS < 0) in both laboratory and wild mice.

## Co-diversified WGM strains out-compete related LGM strains in vivo

The observations that co-diversified LGM strains show genomic evidence of both laboratory-specific adaptation (Fig. 2) and elevated genetic drift (Fig. 3) raised two conflicting hypotheses regarding the relative fitness of LGM and WGM strains. Laboratory-specific adaptation is expected to increase the fitness of LGM strains compared to WGM relatives in the laboratory-mouse environment. In contrast, elevated genetic drift (Fig. 3) is expected to reduce fitness via the accelerated fixation of weakly deleterious mutations (*i.e.*, genetic load). To assess the net consequences of these evolutionary forces in driving divergence between LGM and WGM strains, we analyzed data from competition experiments in which the relative fitness of strains was assessed directly in germ-free laboratory mice^5^.

In these experiments, individual wildling (C57BL/6J harboring wild-derived microbiota), laboratory (C57BL/6J from Jackson Laboratory), and germ-free (GF) (C57BL/6J from Taconic) mice were sampled and co-housed in trios for 17 days^5^. We assessed the relative fitness of co-diversified wildling- and laboratory mouse–specific GM lineages in germ-free mice, enabling comparisons to previous analyses that assessed invasion ability of all GM taxa^5^. To test for the relative fitness of LGM lineages from co-diversified taxa compared to closely related wildling GM lineages, we identified all taxa that showed evidence of co-diversification with hosts (Fig. 1b; Extended Data Table 3) and were present in both wildling and laboratory experimental mice. We then identified all ASVs within these taxa that were detected in either wildling or laboratory mice, but not both, at day 0 (*i.e.*, ASVs that were unambiguously wildling– or laboratory mouse– derived). Based on these ASVs, beta-diversity dissimilarities between wildling and laboratory mice sampled at day-0 (before co-housing) and co-housed GF mice sampled from day 1 to day 17 indicated that wildling GM displayed significant competitive advantages over closely related LGM (Fig. 4). These advantages were evident during the first half of the experiment (days 1–6) and became more pronounced in the latter half of the experiment (days 7–14) (Fig. 4a, b; Extended Data Table 9) (non-parametric permutation t-test *p*-values < 0.001). Longitudinal relative abundances of wildling and laboratory-mouse ASVs within co-diversified genera showed significant wildling advantage (Fig. 4c) (*p*-value < 0.05; t-test, difference in mean abundance days 7–17). Wildling advantage was observed in *Anaerotignum* (*p*-value = 0.027), which also showed genome-wide evidence of elevated genetic drift in the laboratory (Fig. 3d).

**Fig. 4.**
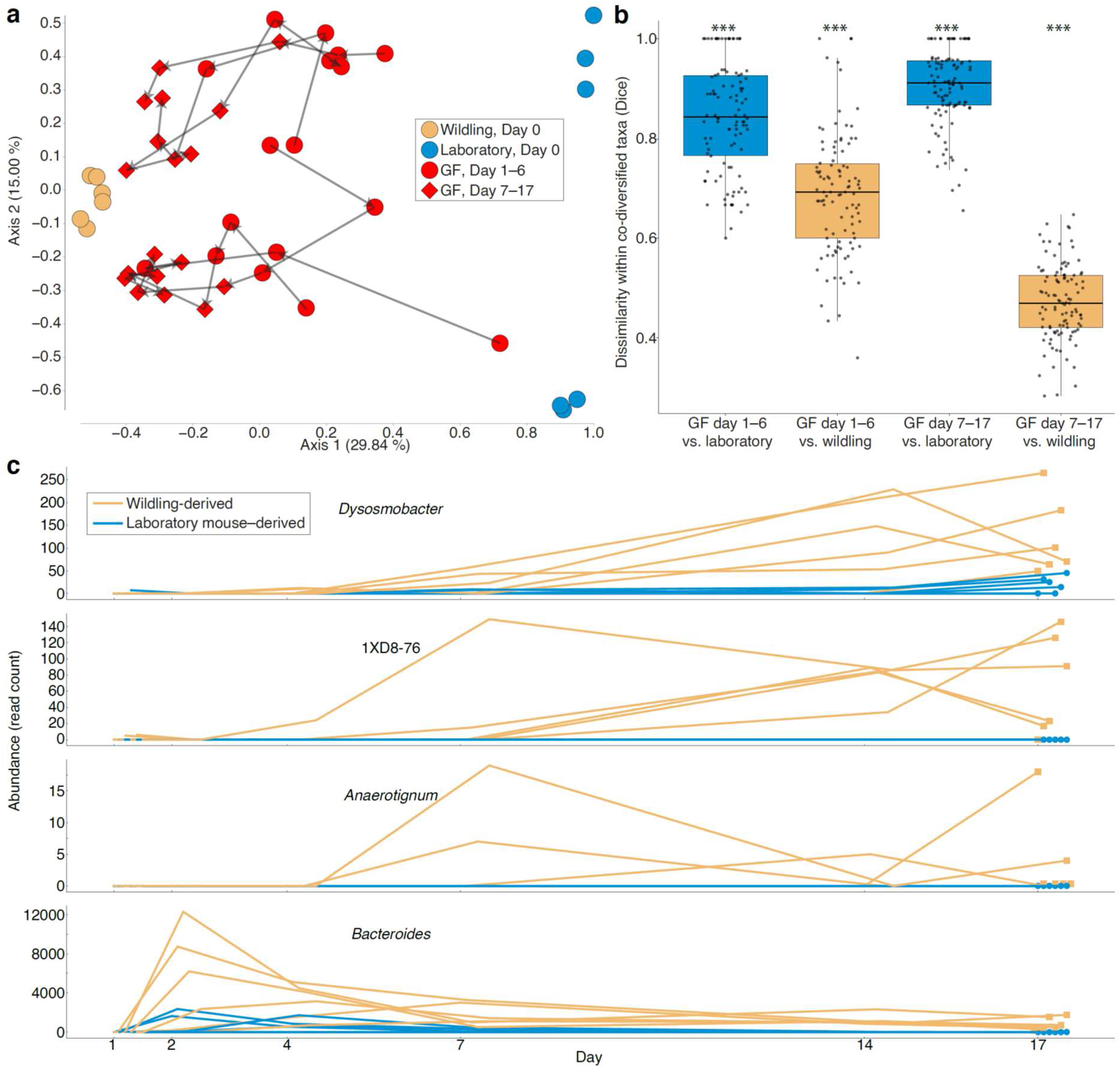
Reduced *in vivo* fitness of co-diversified LGM strains compared to WGM relatives. **a**, Principal coordinates plot shows beta-diversity (Dice) dissimilarities among microbiota profiles based on laboratory mouse– and wildling-specific ASVs within co-diversified taxa. Points are colored based on host of origin as indicated in the key (GF = ex-germ-free mice). Arrows connect longitudinal samples from the same germ-free mouse. **b**, Boxplots show that ex-germ-free mice harbored significantly more wildling-derived ASVs than laboratory mouse– derived ASVs within co-diversified taxa. Asterisks indicate significant differences of boxplot from all other groups based on permutation tests; *** *p*-value < 0.001. Significance testing was conducted after averaging values of longitudinal samples from individual mice within each group. **c**, Line plots show higher abundance of wildling-derived ASVs (yellow) compared to laboratory mouse–derived ASVs (blue) within four bacterial genera showing evidence of co-diversification (Fig. 1). Each line indicates the relative abundance of wildling-derived or laboratory mouse–derived ASVs from each genus within individual ex-germ-free mice between days 1 and 17. Data for other taxa are presented in Extended Data Table 10.

Analyses based on co-diversified taxa indicated stronger advantages for wildling-GM ASVs than did analyses based on the complete dataset containing non-co-diversified taxa (Extended Data Figure 8), and removing co-diversified taxa from the complete dataset reduced the observed wildling competitive advantage (Supplementary Information). These results show that co-diversified GM taxa displayed disproportionate (relative to non-co-diversified GM taxa) advantages for wildling ASVs, supporting accelerated accumulation of genetic load in co-diversified LGM strains.

In summary, we found that laboratory mice have retained > 25-million-year-old symbiont lineages that co-diversified with rodent species, and that these ancestral laboratory-mouse symbionts have experienced elevated levels of genetic drift during > 120 years of captivity. The observation that LGM strains from ancestral, co-diversifying taxa display increased genetic load (Fig. 3) provides an evolutionary basis for their reduced fitness when competed in germ-free mice against relatives from wild mice (Fig. 4). These findings suggest that genetic drift—rather than positive selection—has been the predominant evolutionary force driving divergence of LGM from wild ancestors.

## Methods

### Ethical statement

All procedures conformed to guidelines established by the U.S. National Institutes of Health and have been approved by the Cornell University Institutional Animal Care and Use committee (IACUC: Protocol #2015-0060).

### Sampling of Peromyscus gut microbiota

Samples from *Permoyscus* species sequenced in this study were collected from the University of South Carolina *Peromyscus* Stock Center. Host species sampled included *P. maniculatus bardii* (n=1), *P. leucopus* (n=1), *P. polionotus* (n=2), *P. errimicus* (n=1), *P. californicus* (n=1), and *P. maniculatus sonoriensis* (n=1). Each host lineage was reared in a common laboratory environment on a standard laboratory chow diet. No host lineage was rederived under sterile conditions (*e.g.*, through embryo transfer to a laboratory mouse lineage). These husbandry practices have been previously shown to enable the retention of wild-derived gut microbiota in the laboratory animal-facility environment for > 10 host generations^39^. For sampling, individual rodents were transferred to clean cages and monitored for 1-2 hours, after which fecal samples deposited during that time were collected with sterile tweezers. Fecal samples were immediately placed in empty sterile tubes on dry ice and shipped to Cornell University, where they were stored at -80C until DNA extraction.

### DNA extraction and library preparation

For *Peromyscus* samples sequenced in this study, DNAs were extracted for Nanopore sequencing using a three-step extraction protocol. Steps included 1) osmotic lysis, 2) enzymatic lysis, and 3) bead beating following previously described methods^10^. For each extraction, ∼100mg of starting fecal material was used to ensure sufficient yield for Nanopore sequencing. Libraries were prepared using the Nanopore Ligation Sequencing kit (SQK-LSK110) following the manufacturer-supplied protocols. Separate extractions from the same fecal samples were made for Illumina short-read metagenome sequencing using Qiagen PowerSoil microbiome extraction kits.

### Illumina metagenome sequencing

Libraries for Illumina short-read metagenome sequencing of *Peromyscus* samples were prepared at the Cornell Biotechnology Resource Center (BRC) using their Illumina TruSeq-equivalent ligation library prep protocol (https://www.biotech.cornell.edu/). Libraries were sequenced on an Illumina NovaSeq sequencer at the UC Davis DNA Technologies Core.

### Nanopore sequencing and base calling

Each library was sequenced on the MinION platform using an entire flow cell. Base calling was conducted either in real time or post-sequencing with the Guppy base caller^44^ v3.1.5 using two nVIDIA RTX 3090 Graphical Processing Units (GPUs). The following settings were employed in guppy_basecaller --device “cuda:all” --chunk_size 3000 --chunks_per_runner 768 -- gpu_runners_per_device 4 --qscore_filtering --min_qscore 7 --config dna_r9.4.1_450bps_hac.cfg --calib_detect --compress_fastq --recursive.

### Assembly of Peromyscus MAGs

Contiguous sequences from *Peromyscus* MAGs were assembled from nanopore sequence data and polished with Illumina short-read sequencing data using the snakemake^45^ reticulatus workflow available at https://github.com/SamStudio8/reticulatus. We used the MetaFlye v2.8 ‘spell’ within reticulatus^46^, followed by a polishing pipeline employing four rounds of polishing by Racon^47^, one round of Medaka v1.0.1 [https://github.com/nanoporetech/medaka], and two rounds of Pilon v1.23 polishing with Illumina short-read metagenome data^48^. Finally, contigs likely to be derived from hosts were removed from the polished assembly using the ‘dehumanizer’ step of the reticulatus pipeline against an assembly from the corresponding host species (*P. leucopus*: GCA_004664715.1; *P. polionotus*: GCA_003704135.2; *P. maniculatus*: GCA_003704035.1; *P. californicus*: GCA_007827085.2). Assembled and polished contigs were binned in Anvi’o v6.2 using CONCOCT^49^ and refined manually using anvi-summarize and anvi-refine^50^.

### Phylogenomic analyses

Phylogeny was constructed from the combined set of MAGs from all host species for which > 20 MAGs were available as well as MAGs from *Castor canadensis*, for which only 16 MAGs were available but which represents a basally branching rodent lineage that was not otherwise represented in the data. Core genes from the bac120 collection were identified for each MAG in GTDB-Tk v1.4.1 using the ‘identify’ function^31^. Concatenated core genes were then aligned in GTDB-Tk using the ‘align’ function with default settings^31^. The alignment was then used to infer a phylogenetic tree of the combined set of rodent MAGs in IQTree2 version 2.1.3 using the settings -mset LG,WAG, --seed 0, and -B 1000.

### Scans for co-diversification

To identify co-diversified clades in the rodent MAG phylogeny, we employed an extension of the method developed by Hommola et al.^32^, which uses permutation tests to estimate non-parametric p-values based on the null hypothesis of no association between the symbiont and host evolutionary distances. This workflow yielded a Mantel’s r correlation coefficient for each clade of gut bacteria tested as well as a non-parametric *p*-value indicating the probability of observing by chance a Mantel’s r greater than or equal to that observed in the real data.

Here, we applied these tests to nodes that contained ≥ 3 hosts and ≥7 symbionts, and spanned less than 1/4^th^ of the total bacterial phylogeny, as we reasoned that more deeply diverging clades represent bacterial diversification events that predate the most recent common ancestor of rodents. For each node, the test employed 999 permutations. Only clades with a resulting p-value of < 0.01 and an r coefficient > 0.75 were considered “co-diversifying” for downstream analyses. All code used to conduct these analyses is available in Python at https://github.com/CUMoellerLab/codiv-tools and in R at https://github.com/DanielSprockett/codiv.

In addition, we conducted scans for co-diversification based on dereplicated clades containing only a single MAG per monophyletic clade of MAGs derived from the same host species. For these tests, we randomly selected a MAG from each monophyletic clade and performed PACo^36^, ParaFit^37^, and Hommola’s test^32^ using default settings. All code used for these analyses is available at https://github.com/DanielSprockett/codiv.

### Permutation tests for whether extent of co-diversification exceeds null expectation

In some cases, the MAG clades tested in the co-diversification scan were non-independent due to the underlying tree structure, complicating the adjustment of p-values based on false discovery rate correction. Moreover, in some cases MAGs belonging to the same co-diversifying clade were sampled from multiple individuals per host species, thereby introducing pseudo-replication into tests for co-diversification between host-species lineages. To address these issues and to assess whether there was greater evidence for co-diversification of MAG clades with rodent species in the MAG phylogeny than expected by chance, we conducted additional permutation tests in which the host-species labels were permuted on the host-species phylogeny 100 times and the co-diversification scans were reperformed for each permutation. These analyses yielded a null distribution of the proportion of co-diversifying clades expected to reach significance thresholds (r > 0.75, *p*-value < 0.01) by chance given the underlying structure of and pseudoreplication in the MAG phylogeny. This null distribution was used to calculate a non-parametric *p*-value indicating the probability of observing, by chance under the null hypothesis of no association between bacterial and host evolutionary distances, a number of significantly co-diversifying clades (r > 0.75, *p*-value < 0.01) that was equal to or greater than the number observed in the analyses based on the host phylogeny containing the correct host-species tip labels. All code used to conduct these analyses is available in Python at https://github.com/CUMoellerLab/codiv-tools and in R at https://github.com/DanielSprockett/codiv.

### Molecular clock analyses

We regressed symbiont divergence estimates based on protein-sequence divergence in clades that displayed the strongest evidence of co-diversification (that is, Mantel’s r > 0.95) and known divergence times of host species based on timetree.org^29^. These regression analyses and calculations of 95% confidence intervals were conducted in base R (version 4.2.3).

### Phylogenetic ANOVA

Phylogenetic ANOVA was performed using the rodent gut bacterial phylogeny and the KEGG annotations for each MAG to identify gene annotations enriched in co-diversifying clades relative to non-co-diversifying clades or in laboratory-mouse symbionts relative to wild-mouse symbionts. Theses analyses were conducted using the phylANOVA function in the R package ‘phytools’^51^ v2.1. Benjamin-Hochberg correction was performed to account for multiple testing across classes of annotations.

### dN/dS analyses

For each co-diversifying clade containing MAGs from laboratory and wild house mice, we used Roary^52^ v3.12.0 to identify and codon-align each gene family containing orthologs from at least one laboratory-derived house-mouse MAG and at least one wild-derived house-mouse MAG. Codon alignments were then used to construct a phylogenetic tree for each gene family using RAxM^53^ v8.2.12. Codon alignments and gene trees were then used in CODEML within PAML^54^ v4.10.6 to estimate the proportion of nonsynonymous substitutions per nonsynonymous site (dN) to synonymous substitutions per synonymous site (dS) (*i.e.*, dN/dS) for every branch leading to a laboratory-derived house-mouse gene copy and every branch leading to a wild-derived house-mouse gene copy. The averages for laboratory-derived house-mouse gene copies and wild-derived house-mouse gene copies were calculated for each gene family. Differences in genome-wide dN/dS across the whole dataset and in individual clades were conducted with paired t-tests in base R. All genes with nonzero dN and dS are shown in Figs. 2 and 3.

### Reanalysis of co-housing experiments

We downloaded fastq files from accessions from PRJNA540893^5^, which contained results of an experiment in which C57BL/6J mice from the Jackson Laboratory (a source from which genomes of laboratory-mouse GM strains displaying evidence of genetic drift were assembled, *e.g.*, Fig. 3) containing either a laboratory-derived or wild-derived microbiota were co-housed with germ-free mice for 17 days. Sequences were denoised with dada2^55^ in qiime2^56^ v2023.9 using the following settings: --p-trim-left 0 --p-trunc-len 200. GTDB ribosomal sequences and taxonomy (bac120_ssu_reps.fna.gz and bac120_taxonomy.tsv) were imported into qiime using ‘qiime tools import’, and a classifier was trained using ‘qiime feature-classifier fit-classifier-naive-bayes’^57^. ASVs were then classified using ‘qiime feature-classifier classify-sklearn’^57^. Samples were rarefied to a common depth of 40,000 reads (results were qualitatively identical with and without rarefaction). A total of 2,640 ASVs were present in the final complete dataset. Diagnostic ASVs belonging to co-diversifying taxa and found in either wildling or laboratory mice at experimental day 0 (105 ASVs) were retained using ‘qiime feature-table filter-features’. Wildling and laboratory-mouse samples at day zero and GF-mouse samples from days 1–17 were retained with ‘qiime feature-table filter-samples’. Dice dissimilarities (to assess strain sharing) among diagnostic ASV profiles were calculated with ‘qiime diversity beta’. Principal Coordinates Analyses (PCoA) were conducted using ‘qiime diversity pcoa’ and plots were generated using ‘qiime emperor plot’^58^. Boxplots were created in R using ggplot2 (Version 3.5.1).

## Acknowledgements

We thank Dr. Weiwei Yan for assistance with DNA extractions from rodent fecal samples. We thank the staff at Peromyscus Stock Center for providing fecal samples.

## Author information

### Contributions

A.H.M. supervised the research, performed analyses, and wrote and edited the paper. D.D.S. performed analyses and wrote and edited the paper. A.A.L., B.A.D., and J.G.S. performed analyses and edited the paper.

## Corresponding author

Correspondence to Andrew H. Moeller.

## Data availability

All sequence data generated in this study have been deposited to the National Center for Biotechnology Information Sequence Read Archive under accessions BioProject ID PRJNA1089132. Additional metadata about the genome assemblies generated by previous studies are available at doi.org/10.1038/s43705-021-00053-9.

---

**Extended Data Figure 1:**
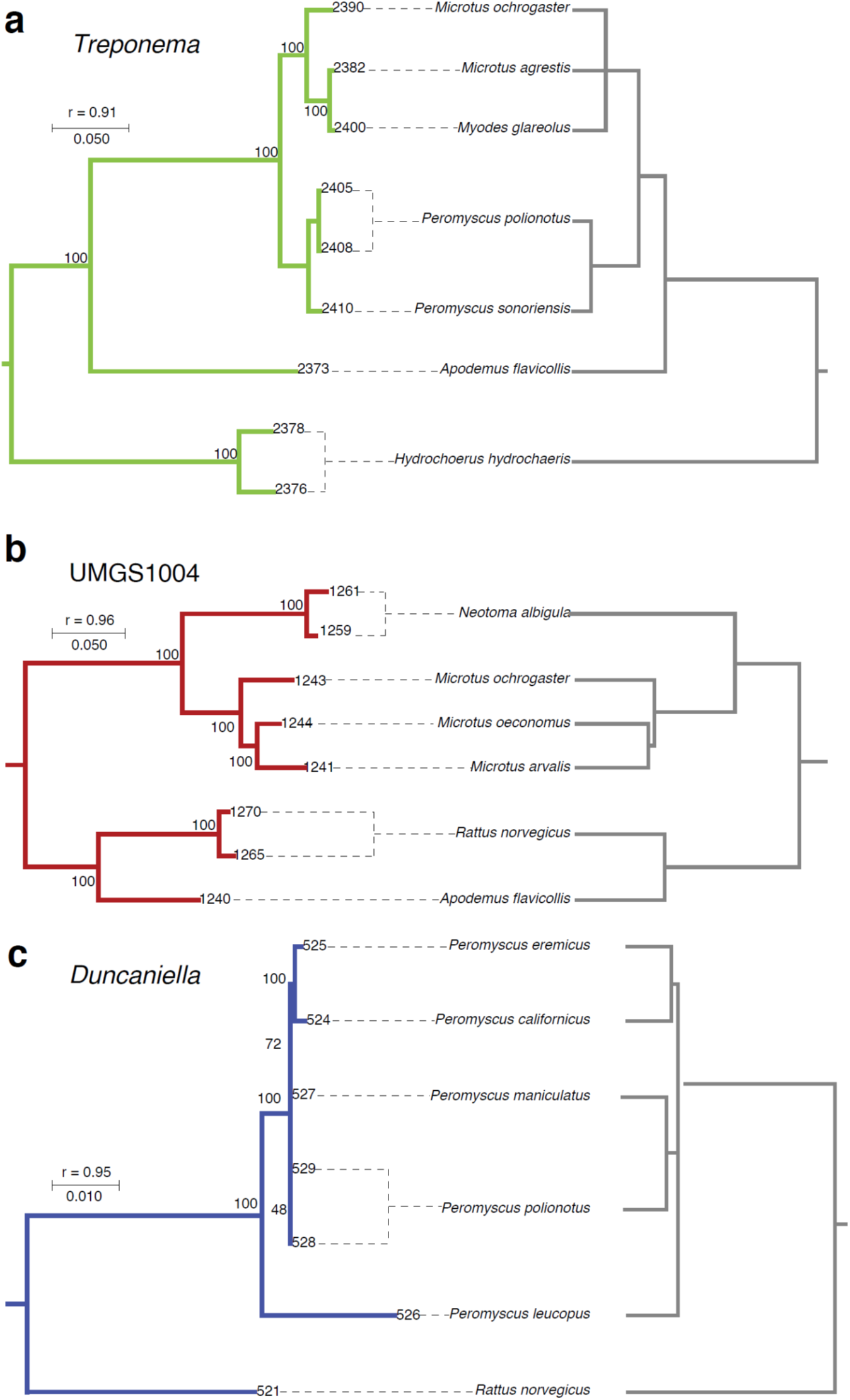
Examples of co-diversifying clades lacking MAGs from house mice. **a–c**, Trees show phylogenetic relationships among symbiont MAGs (left) and the rodent host species from which they were recovered (right). Symbiont branches are colored by phylum as in Fig. 1. Dashed lines connect MAGs to the host species from which they were recovered. The test statistic from the Hommola test (r) for each clade is shown. Branch lengths correspond to amino acid substitutions per site.

**Extended Data Figure 2:**
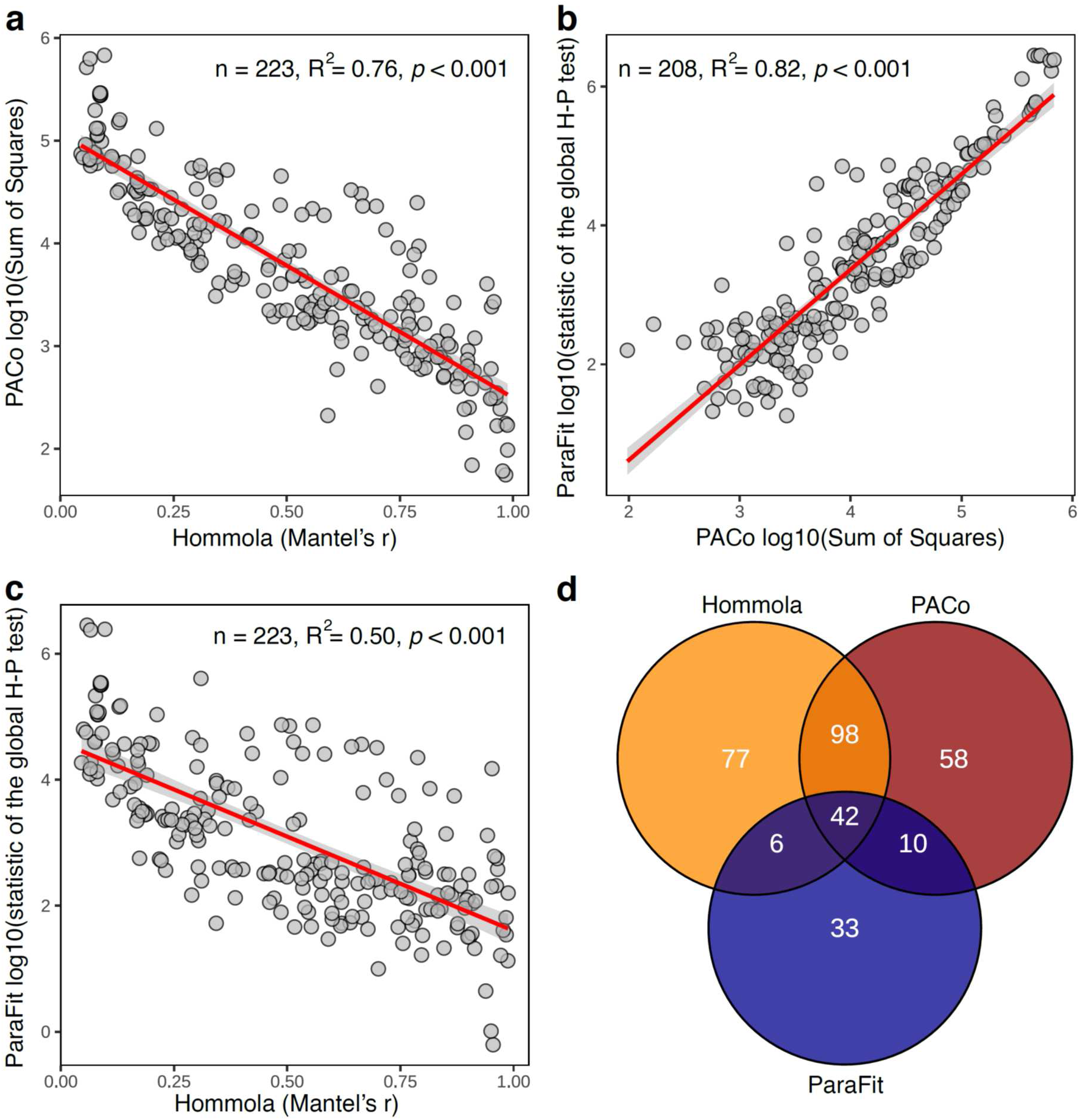
Comparisons of tests for co-diversification based on de-replicated (*i.e.*, collapsed) tests. **a–c**, Scatter plots show relationships of test statistics between pairs of tests for co-diversification based on clades subsampled to a single MAG per host species. Tests include Hommola, PACo, and ParaFit. Red lines indicate best-fit regressions, and shaded gray areas represent 95% confidence bands. **d,** Venn diagram shows overlap between significant clades obtained from the three different tests employed.

**Extended Data Figure 3:**
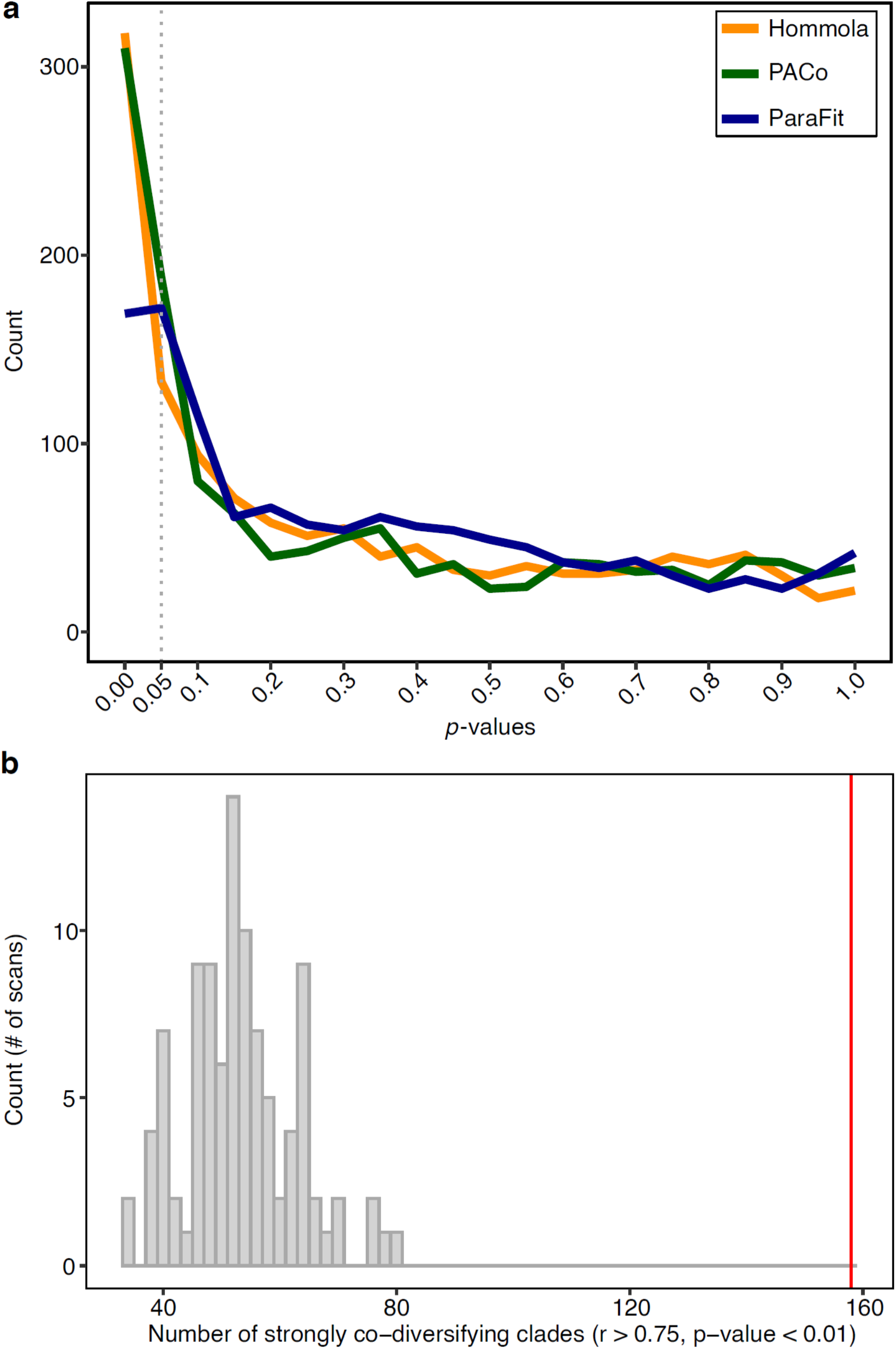
Evidence for co-diversification from each class of tests exceeds that expected under the null hypotheses. **a**, Lines show distributions of *p*-values obtained from scans of co-diversification based on down-sampled clades (*i.e.*, down sampled to contain one MAG per host species) for PACo, ParaFit, and Hommola tests. **b,** Histogram shows the number of co-diversifying clades detected (r > 0.75, *p*-value < 0.01) based on Hommola scans of the entire MAG phylogeny and permuted host-species tip labels. Vertical red line indicates the number of co-diversifying clades detected in the scan based on the real data (*i.e.*, non-permuted host-species tip labels).

**Extended Data Figure 4:**
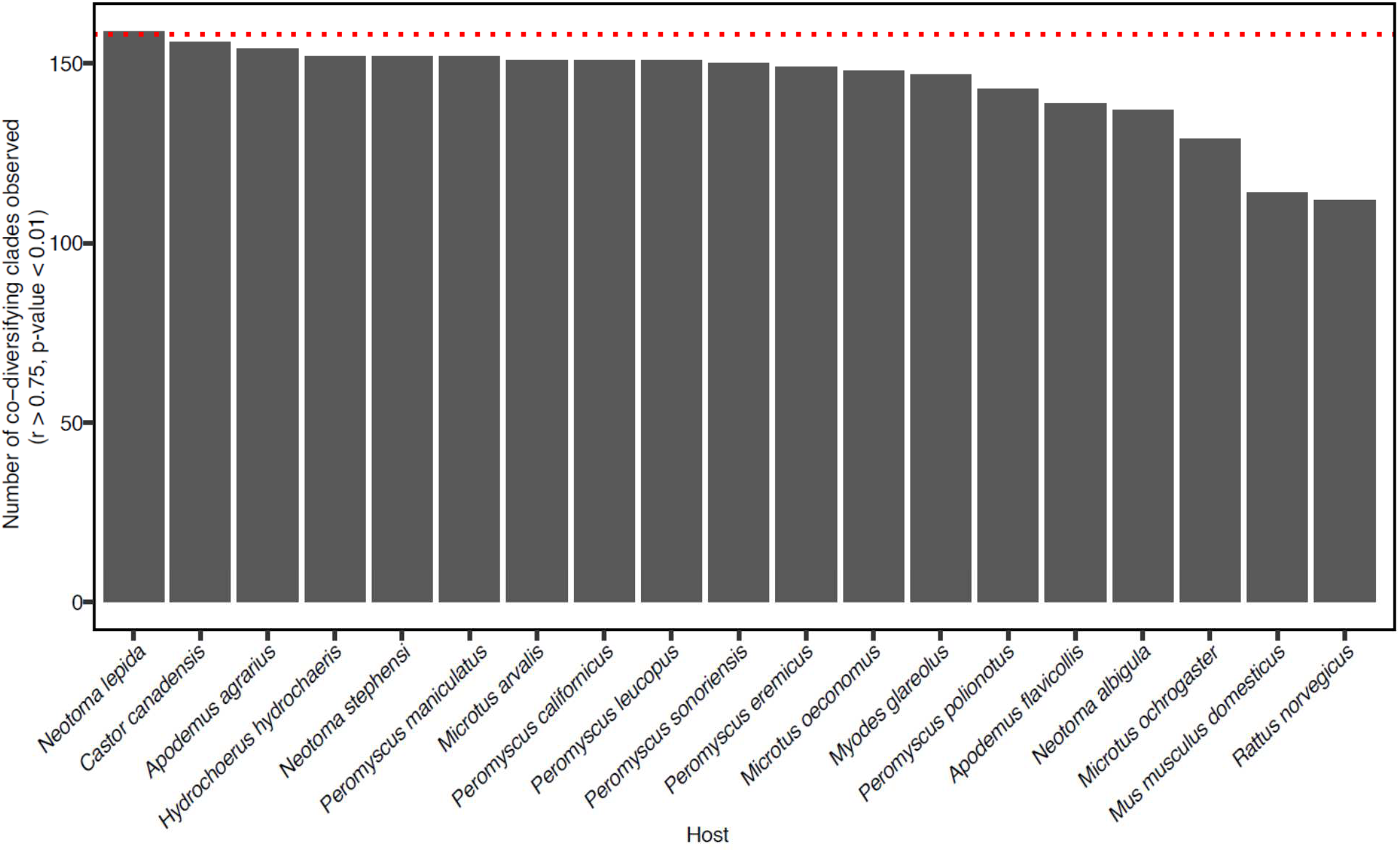
Sensitivity analyses of Hommola scans for co-diversification based on the complete dataset. Barplot shows the number of strongly co-diversifying clades (r > 0.75, *p*-value < 0.01) detected based on sensitivity analyses in which scans for co-diversification were performed after removing each host species one at a time. X-axis shows which host species was removed from the scan.

**Extended Data Figure 5:**
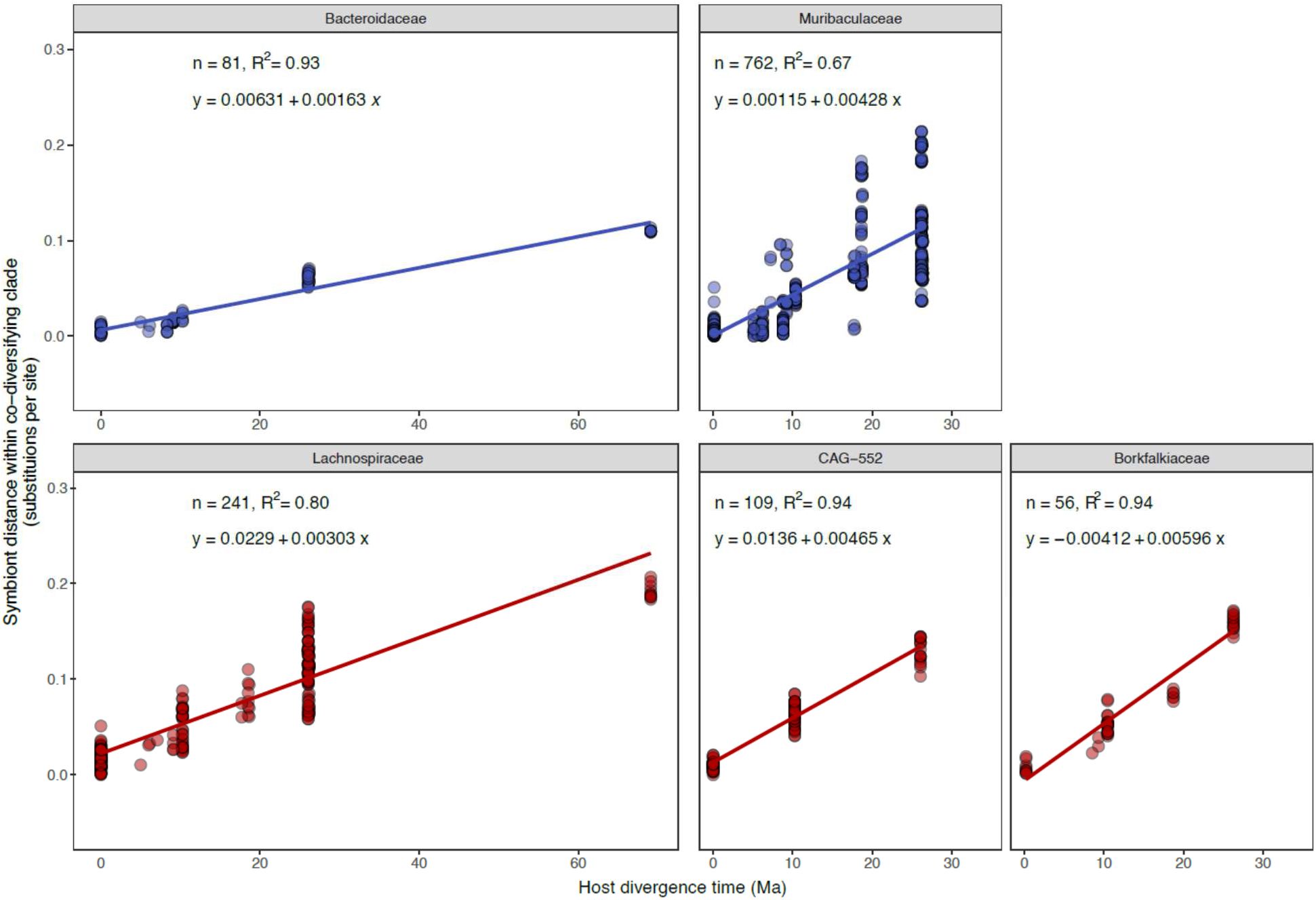
Molecular clock analyses corroborate co-diversification. (A) Scatter plots show the relationship between protein sequence divergence within strongly co-diversifying symbiont clades (Mantel’s r > 0.95) (y-axis) and the ages of divergence events between host clades from which they were recovered (x-axis). Lines show best-fit regressions between pairwise symbiont distances within co-diversifying clades and host divergence times. Plots are colored based on bacterial phylum as in Fig. 1, and all families containing > 2 co-diversifying clades (based on non-downsampled Hommola tests) are shown.

**Extended Data Figure 6:**
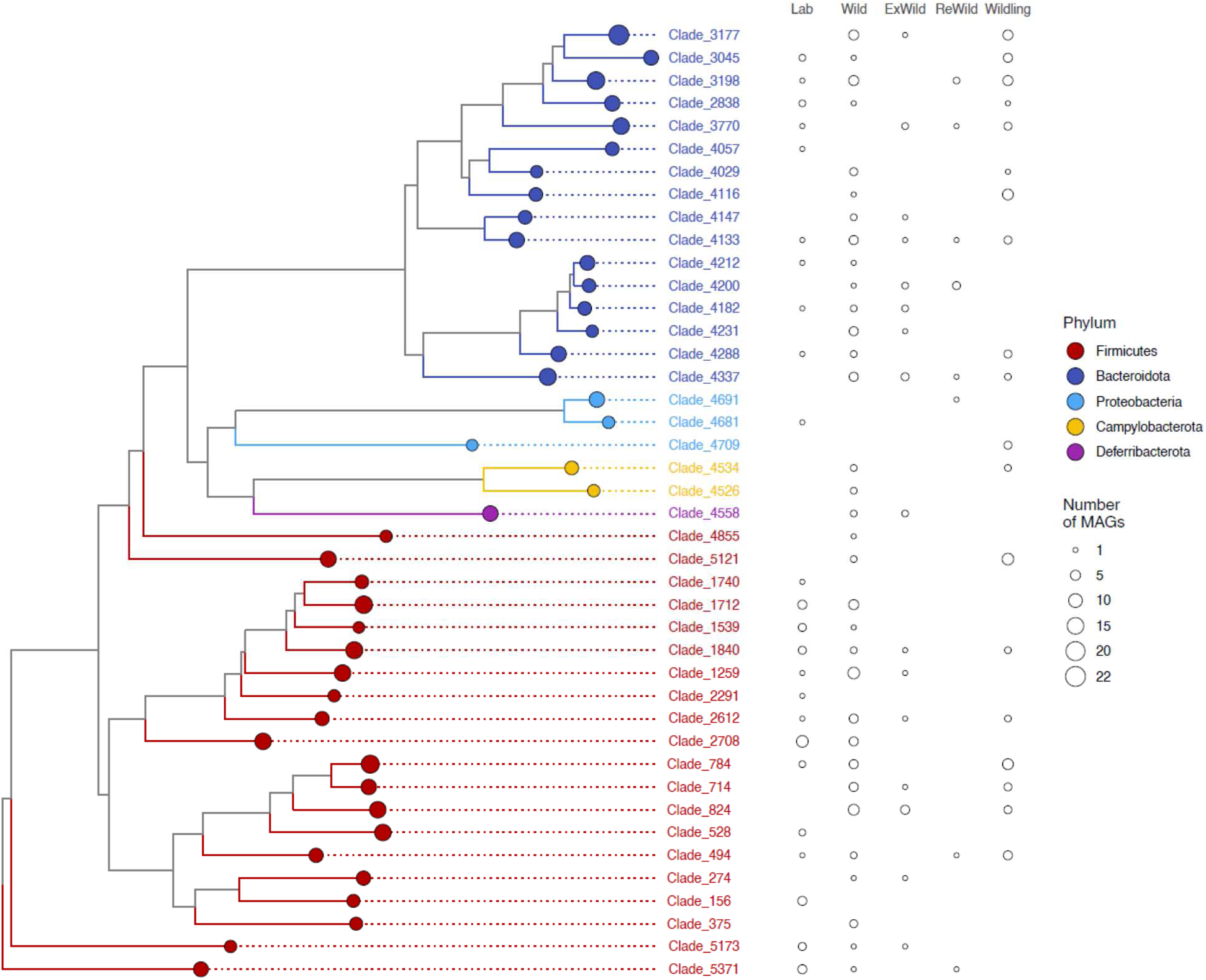
Retention and extinction of ancestral, co-diversifying symbioses from laboratory house mice. Phylogeny shows relationships among all bacterial genomes belonging to co-diversifying clades shown in Fig. 1 that contain genomes from house mice, at least one other murid host, and at least one non-murid host, *i.e.*, co-diversifying clades that could be inferred to be ancestral to murids (the rodent family containing house mice). Colored circles indicate clades. Sizes of circles indicate the number of MAGs contained in the clade. Circles to the right of the phylogeny indicate the number of MAGs in each clade derived from wild mice, laboratory mice, ‘rewilded’ mice (*i.e.*, lab born but released and sampled outdoors), ‘ex-wild’ mice (wild-caught mice brought into the lab), or wildling mice (*i.e.*, laboratory genotype born to wild-caught mother via embryo transfer). Rightmost two columns highlight how co-diversifying clades that have been lost from laboratory mice can be regained in rewilded or wildling mice.

**Extended Data Figure 7:**
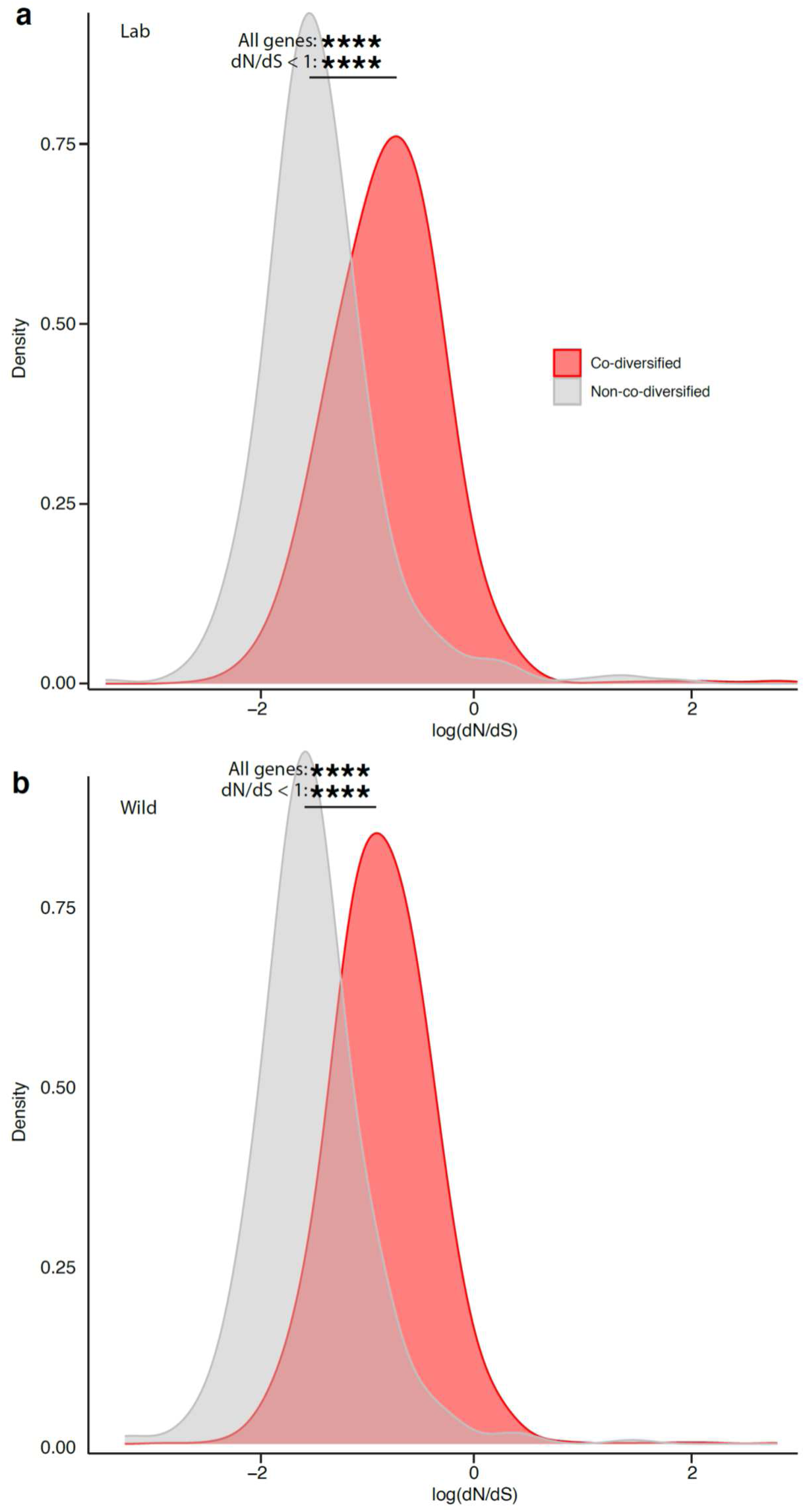
Significantly elevated genome-wide genetic drift in co-diversifying relative to non-co-diversifying GM strains. **a, b**, Density plots show significant genome-wide elevation of dN/dS (a hallmark of genetic drift) in co-diversifying versus non-co-diversifying GM clades in both laboratory house mice (**a**) and wild house mice (**b**). Significance of t-tests for differences in mean are denoted by asterisks; **** *p*-value < 0.0001. Top asterisks denote significance of tests considering all genes, and bottom asterisks denote significance of tests considering only genes displaying dN/dS < 1 (log dN/dS < 0) in both laboratory and wild mice.

**Extended Data Figure 8:**
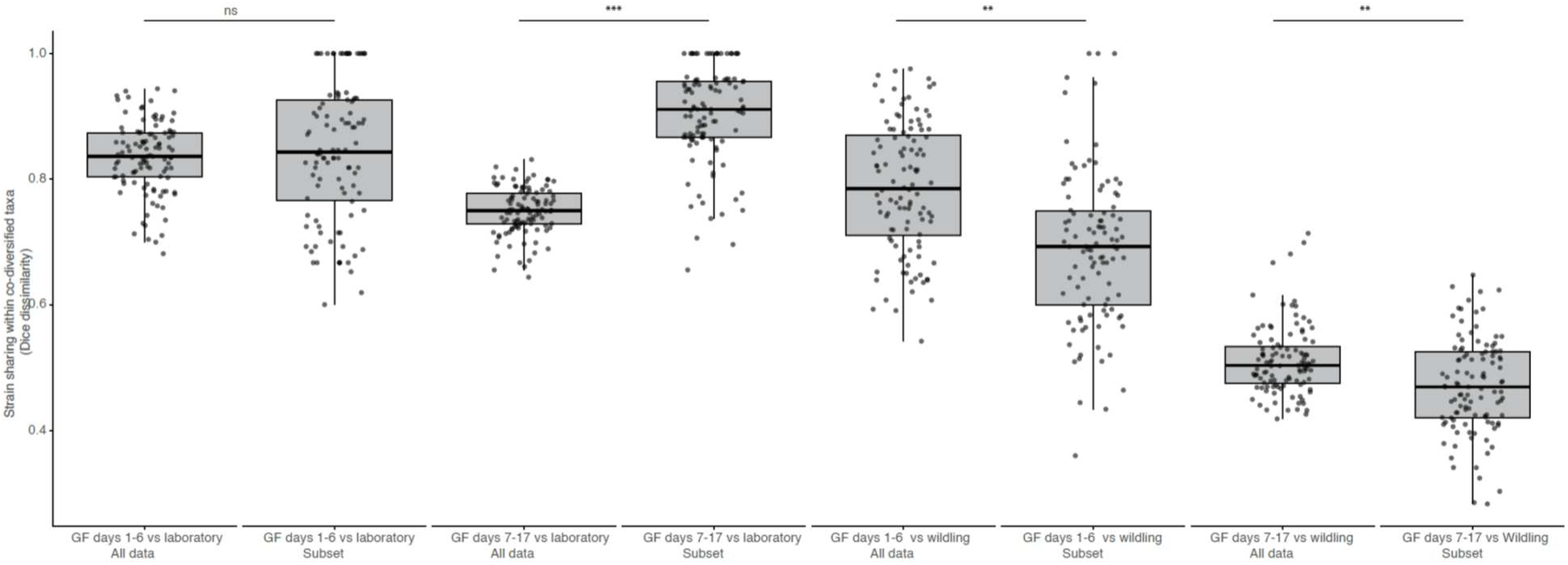
Disproportionate competitive advantage of wildling ASVs from co-diversified taxa. Boxplots show microbiota dissimilarity (Dice) based on the complete dataset (All data) or only co-diversified ASVs retained for the analyses whose results are shown in Fig. 4. Asterisks indicate significant differences based on permutation t-tests; *** *p*-value < 0.001, ** *p*-value < 0.01. significance testing was conducted after averaging values of longitudinal samples from the individual mice within each group. Note the increased dissimilarity between GF and laboratory microbiota in results based on subsetted ASVs (right box within each facet) compared analyses based on to all ASVs (left box within each facet) (particularly from days 7–14). Similarly, note the decreased dissimilarity between GF and wildling microbiota in results based on subsetted ASVs compared analyses based on to all ASVs.

**Extended Data Table 1. Metadata for *Peromyscus* samples sequenced with Nanopore.**

**Extended Data Table 2. Metadata for MAGs.**

**Extended Data Table 3. Results from co-diversification tests.** Columns labeled as ‘collapsed’ show results of tests of clades down sampled to a single MAG per host species. Note that the last 25 rows do not contain symbiont trees due to string-length limits in Microsoft Excel. The Newick strings for these subclades (none of which showed significant evidence of co-diversification) are available at https://github.com/DanielSprockett/codiv.

**Extended Data Table 4. Pairwise comparisons for molecular clock estimates within co-diversifying clades.**

**Extended Data Table 5. Co-diversified (r > 0.75) symbiont gene families under positive or purifying selection in laboratory or wild house mice.**

**Extended Data Table 6. Gene families enriched in laboratory or wild house mice based on phylogenetic ANOVA.**

**Extended Data Table 7. Gene families enriched in co-diversifying or non-co-diversifying clades based on phylogenetic ANOVA.**

**Extended Data Table 8. Non-co-diversified (r < 0) symbiont gene families under positive or purifying selection in laboratory or wild house mice.**

**Extended Data Table 9. Dice dissimilarities between samples from Rosshart et al.^13^ based on laboratory- and wild-specific ASVs within co-diversified taxa.**

**Extended Data Table 10. Read counts for laboratory- and wildling-specific ASVs from co-diversified taxa in samples from ex-germ-free mice.**

## Supplementary Information

### Nanopore metagenomic sequencing of wild-derived Peromyscus lineages

We deeply sequenced the metagenomes of six species/subspecies of *Peromyscus*, including, sampled at the *Peromyscus* stock center at the University of South Carolina, Columbia. Seven MinION flow cells were used to sequence DNAs extracted from fecal samples, with one flow cell dedicated to fecal samples from each individual host. The final *Peromyscus* dataset contained 30,612,212 reads, ranging from 2,325,236 to 6,018,007 reads per sample. The average per-sample read length ranged from 3,494 to 8,335 base pairs. Metadata for all *Peromyscus* samples analyzed in this study are presented in Extended Data Table 1.

### Long-read assembly of bacterial genomes from Peromyscus metagenomes

Assembling and binning contigs from long-read metagenomes generated by Nanopore sequencing of fecal samples from the *Peromyscus* species yielded a total 504 metagenome assembled genomes (MAGs) of high-quality (> 50% completeness < 5% contamination) from the six host species. Bacterial diversity represented in these MAGs spanned 10 phyla, including Actinobacteriota, Bacteriodota, Campylobacterota, Deferribacterota, Desulfobacterota, Firmicutes, Patescibacteria, Proteobacteria, Spirochaetota, and Verrucomicrobiota. Taxonomic assignments of all MAGs newly generated by this study are presented in Extended Data Table 2.

### Phylogenetic analyses of rodent MAGs and hosts

Single copy bac120 core genes from each MAG were identified and aligned with GTDB-Tk, and the alignment was used for phylogenetic inference with IQTree2 v2.2.0.4 with the following parameters: --seed 0 -B 1000 -alrt 1000 -mset WAG,LG. *P. maniculatus sonoreinsis* was not available in the timetree.org database and was therefore placed manually in the host phylogeny as sister to *P. maniculatus bardii* with a divergence time of 500,000 years.

### Sensitivity analyses of co-diversification results

We performed a sensitivity analysis to assess the impact of the MAGs from each individual host species by conducting the Hommola co-diversification scan on each possible subset of MAGs containing all MAGs except those from an individual host species. This analysis tested whether the results observed in Fig. 1 depended on MAGs from any individual host species. The results show that the detection of most co-diversifying clades was robust to the exclusion of MAGs from any individual host species (Extended Data Figure 4). The exclusion of MAGs from *Mus musculus domesticus* or *Rattus norvegicus*—the two host species represented by the most MAGs—had the largest impact on the number of co-diversifying clades detected. However, even when MAGs from one of these host species were excluded, scans identified > 100 co-diversifying clades.

*Calibration of molecular clocks in the rodent gut microbiota corroborates co-diversification* Given the phylogenetic evidence that symbiont lineages and host species co-diversified, the known divergence dates of host species based on molecular data and fossils can be used to calibrate bacterial molecular clocks^59–61^, which are otherwise difficult to calibrate due to the lack of a bacterial fossil record. Symbiont and host evolutionary distances within co-diversifying clades were positively associated in all bacterial families (Families containing > 2 co-diversifying clades are shown in Extended Data Figure 5. All data are presented in Extended Data Table 4), enabling calibration of the rates of molecular evolution in diverse GM taxa. Rates ranged from 0.00163 to 0.00596 substitutions per million years. These rates are within the range estimated previously from codiversifying symbionts in primates^10^ and timeseries data of bacterial pathogens^62–64^, further supporting the concurrent diversification of bacterial and rodent host lineages.

### Identification of clades ancestral to Muridae

We identified all co-diversifying clades that contained representatives from house mice, a non-house mouse murid and at least one outgroup to the Muridae (40 non-nested, *i.e.*, independent, clades). Identifying these clades allowed us to generate a set of clades ancestral to Muridae independent of data from house mice, thereby enabling us to assess rates of extinction and retention of these clades from either laboratory or wild house mice using a common set of ancestral co-diversifying clades.

*Differentially abundant gene families between co-diversifying and non-co-diversifying clades* To identify gene families enriched or depleted in co-diversifying rodent symbionts relative to non-codiversifying rodent gut bacteria independent of bacterial phylogenetic history, we annotated each MAG using the Kyoto Encyclopedia of Genes and Genomes (KEGG) ontogeny. Next, we employed phylogenetic ANOVA^51^ using the rodent gut bacterial phylogeny to identify annotations over- or under-represented in co-diversifying gut bacteria (r > 0.75, *p*-value < 0.01) relative to non-co-diversifying gut bacteria (r < 0). No individual annotation reached significance after correction for multiple testing. These analyses provided a rank order list of gene annotations significantly associated (positively or negatively) with co-diversification (Extended Data Table 7).

### Tests for gain and loss of functions in genomes of laboratory–house mouse symbionts

In addition to adaptive evolution of protein sequences, gut bacterial genomes can adapt to novel environments through changes in gene content. Genes that benefit fitness in the new environment can be gained by gene duplication or horizontal transfer and favored in bacterial populations by positive selection, whereas ancestral genes that no longer provide appreciable fitness benefits can be deleted by mutation (which displays a bias towards deletion in bacteria^65,66^) and lost from bacterial populations by genetic drift (or, for costly genes, by negative selection). To test whether the genomes of co-diversified symbionts have experienced laboratory-specific expansions or contractions of specific gene families, we performed phylogenetic ANOVA for each gene family in each co-diversifying symbiont clade containing MAGs from both wild and laboratory mice (Supplementary Information). These analyses asked whether the genomes of multiple, phylogenetically independent symbiont lineages have gained or lost—convergently or in parallel—the same gene families in response to the transition from the wild into captivity. These analyses provided a rank-order list of genes enriched or depleted in MAGs from laboratory house mice relative to wild house mice. No gene functions were significantly enriched or depleted after false-discovery correction.

### Lack of elevated genetic drift in non-co-diversifying GM clades

In addition to testing for elevated genetic drift in laboratory-mouse GM strains in co-diversifying clades (r > 0.75), we also tested for elevated genetic drift in laboratory-mouse GM strains in non-co-diversifying clades (r < 0) of similar phylogenetic depth to co-diversifying clades (*i.e.*, clades of congeneric MAGs in the distal 1/4^th^ of bacterial phylogeny). These analyses did not support a significantly increased genetic drift, as measured by genome-wide elevation of dN/dS, in non-codiversifying clades (paired t-test p-values = 0.0944 for tests based on genes under purifying selection, *i.e.*, genes for which dN/dS was < 1 in both wild-mouse and lab-mouse GM strains), contrasting the results observed for co-diversifying clades (Extended Data Table 8). Tests on individual non-co-diversifying clades also failed to strongly support increased genome-wide dN/dS in non-co-diversifying laboratory-mouse GM strains (paired t-test FDR-corrected p-values > 0.01 for all clades, and > 0.1 for all but 1 clade). A single clade (node 5493: an unclassified genus, CAG-1435, in the order Christensenellales) showed a marginally significant elevation in dN/dS in laboratory-mouse GM strains relative to WGM strains (paired t-test FDR-corrected p-value = 0.0241), and when all genes were tested, a marginally significant increase in dN/dS in the laboratory-mouse GM strains was observed (paired t-test p-values = 0.04065). This latter difference can be attributed to genes showing evidence of positive selection (dN/dS > 1) in laboratory-mouse GM strains but purifying selection (dN/dS < 1) in WGM strains, rather than increased genetic drift. Cumulatively, these results indicate that the significant elevation of genome-wide dN/dS (indicative of reduced *Ne* and increased genetic drift) in laboratory house mice observed for co-diversified GM strains was not apparent for non-co-diversified GM strains.

### Elevated genetic drift in co-diversifying relative to non-co-diversifying clades in both laboratory and wild house mice

Significant evidence of elevated genetic drift in laboratory GM strains was detected in co-diversifying clades but not non-co-diversifying clades (Tables S5, S8), suggesting that co-diversifying clades may be particularly predisposed to elevated genetic drift. Previous studies of insect endosymbionts have shown that bottlenecks during transmission of host-restricted symbionts can promote genetic drift^12,13^, but the extent to which host-restriction of GM symbionts in mammals promotes genetic drift has not been explored. To address this idea, we tested whether co-diversifying clades displayed stronger evidence of genetic drift than non-co-diversifying clades regardless of environment (laboratory or wild). We compared the distributions of per-gene log(dN/dS) values between co-diversified and non-co-diversified clades in both the laboratory and the wild. Results indicated significant elevation of genetic drift, as indicated by elevated genome-wide dN/dS, in co-diversified relative to non-co-diversified GM clades (Extended Data Figure 7) in both the laboratory (Extended Data Figure 7A) and the wild (Extended Data Figure 7B). These findings suggest that the host restriction of co-diversified clades predisposes these lineages to stronger genetic drift, which can be further enhanced by transitions from the wild to the laboratory environment (Fig. 3).

### Removal of co-diversifying ASVs reduces signal of competitive advantage for wildling microbiota

To test whether ASVs of wildling or laboratory origin within co-diversifying taxa displayed a disproportionately strong competitive differential (compared to all ASVs), as suggested by Extended Data Figure 8, we compared results of beta-diversity analyses based on the complete dataset with those based on the complete dataset minus the ASVs used in analyses whose results are shown in Fig. 4. This comparison allowed us to test whether the removal of these ASVs belonging to co-diversifying taxa reduced the measured competitive differential between wildling and laboratory microbiota, as expected if co-diversifying laboratory GM strains have acquired increased genetic load. Indeed, removing these ASVs led to weaker signal of competitive advantage for wildling microbiota. On average, when all ASVs were included, the Dice beta-diversity differential between GF day 7–14 versus laboratory microbiota and GF 7–14 versus wildling microbiota was 0.237 (favoring wildling microbiota), whereas when only the subsetted ASVs (as defined in Fig. 3 and Extended Data Figure 8) were included this differential reduced to 0.22 (the difference in differential of only ∼0.01, given that the subsetted ASVs constituted 105 of 2640 total ASVs. These findings indicate that laboratory-derived GM strains belonging to co-diversifying taxa displayed disproportionately stronger evidence for reduced fitness in these experiments than did other laboratory-derived GM strains, mirroring results presented in Extended Data Figure 8. These findings further support increased genetic load in laboratory-derived GM strains in co-diversifying taxa.

